# Are Deep Learning Structural Models Sufficiently Accurate for Free Energy Calculations? Application of FEP+ to AlphaFold2 Predicted Structures

**DOI:** 10.1101/2022.08.16.504122

**Authors:** Thijs Beuming, Helena Martín, Anna M. Díaz-Rovira, Lucía Díaz, Victor Guallar, Soumya S. Ray

## Abstract

The availability of AlphaFold2 has led to great excitement in the scientific community - particularly among drug hunters - due to the ability of the algorithm to predict protein structures with high accuracy. However, beyond globally accurate protein structure prediction, it remains to be determined whether ligand binding sites are predicted with sufficient accuracy in these structures to be useful in supporting computationally driven drug discovery programs. We explored this question by performing free energy perturbation (FEP) calculations on a set of well-studied protein-ligand complexes, where AlphaFold2 predictions were performed by removing all templates with >30% identity to the target protein from the training set. We observed that in most cases, the ΔΔG values for ligand transformations calculated with FEP, using these prospective AlphaFold2 structures, were comparable in accuracy to the corresponding calculations previously carried out using X-ray structures. We conclude that under the right circumstances, AlphaFold2 modeled structures are accurate enough to be used by physics-based methods such as FEP, in typical lead optimization stages of a drug discovery program.

## Introduction

Despite progress in structural biology, including the advent of novel cryo-EM methods^1^, experimental structures remain unsolved for a large portion of druggable targets in the human genome^2^. During the past year, however, new developments in deep learning approaches have revolutionized the world of structural biology. For the first time, drug discovery projects can leverage the use of structural data in cases where experimentally resolved structures (or those of very close homologs) are not available. This is made possible by the pioneering work from DeepMind, who recently developed and released the AlphaFold2 (AF2) code^3^. The AF2 methodology, along with similar techniques^4^, has shown unprecedented results when predicting structures from sequence alone, leading to a dramatic increase in accuracy, and potentially widening the domain of applicability of structure-based design. In these methods, models are built by using physics-based and knowledge-based energy functions, combined with evolutionary information (at a pair representation level) enabling spatial and evolutionary relationships. This has enabled genome wide application of structure prediction, resulting, for example, in the availability of a structural database for the human proteome^5^.

Following the open-source release of the AF2 code, several studies have modified the original algorithm and have attempted to determine the applicability of deep learning methods to a range of structural problems, including the identification and characterization of protein-protein interactions ^6,7^, the prediction of protein-peptide complexes^8^, and the modeling of conformational transitions for drug receptors^9^. Beyond global structural and fold prediction, there is an obvious need to determine whether structures predicted with AF2 (or related methods) are sufficiently accurate for use in in silico screening or hit-to-lead modeling, especially in situations where there is limited structural information available (i.e., no availability of close structural or sequence homologs). Besides this, given that AF2 relies on an exhaustive and elaborate training process, it is important to understand the effects of the presence of closely related homologs of query sequences in the original training sets on the conformations of the resulting models.

In parallel, in the last few years we have witnessed important advances in computational chemistry methods which, together with the dramatic exponential growth in computational power, have led to an increased application of structure-based design in drug discovery projects. Physics-based computational approaches are now routinely used to predict a range of properties, from potency to solubility, at various stages of the drug discovery pipeline, including lead identification and lead optimization^10^. In particular, recent advances in force fields and sampling algorithms have now made it possible to use free energy methods to calculate relative affinities of compounds for proteins with accuracies of < ~1.0 kcal/mol^11^, which starts to approach the experimental accuracy of most biochemical and biophysical assays for protein-ligand interactions^12^. The increased accuracy of computational methodologies indicates that the domain of applicability of structurebased approaches is now largely limited by the availability of a high-resolution structure of a ligand-protein complex.

It remains an open question whether AF2 models for novel protein folds (meaning structure for which no close structural homologs are available) are accurate enough for physics-based prediction methods, including computational approaches such as virtual screening and free energy calculations that require understanding of the details of the protein – ligand complex. To address this, we assessed whether a physics-based method for predicting compound potency (Free Energy Perturbation, or FEP) can be successfully used in combination with ab-initio models developed using AF2. We have applied a best-in-class implementation of FEP (Schrödinger’s FEP+^13^) to a series of AF2 modeled targets (details in methods section), where its accuracy has already been demonstrated when applied to crystal structures^11,14,15,16^, making a direct assessment of the relative performance of these AF2 models possible. In addition, we performed this experiment by simulating a scenario where no template structures with high sequence identity (>30%) were available for developing accurate homology models.

To this end, we have developed a custom version of AF2 where we systematically removed template structures and homologous sequences from the database, aiming at reproducing a situation where traditional homology model techniques have been shown to fail, for example in blinded prospective tests such as CASP^17,18^. Our results demonstrate that in a realistic prospective scenario, with only homologs of less than 30% sequence identity available, AF2 is capable of accurately providing structural models and, more importantly, can be used to predict relative changes in ligand affinities with an accuracy that is statistically comparable to those obtained using crystal structures.

## Results

### Dataset selection

In order to test whether AF2 models were suitable as starting points for running state-of-the-art Free Energy Perturbation calculations, we tried to reproduce affinity predictions previously obtained in benchmarks using the Schrödinger implementation of FEP (FEP+^13^). We assembled a number of datasets that were part of these prior FEP+ benchmarks (see Table 1), including 8 targets studied in the original description of the method^11^, two targets obtained from a benchmark dedicated to fragments^14^, two targets studied from a study of application of FEP G-protein-coupled receptors^15^, and two targets from a publication describing application to selectivity studies^16^.

**Table 1.** Global and binding site Cα-atom RMSD of models produced with AF2, aligned with the reference PDB. The binding site includes all residues within 5Å from the ligand in the reference PDB. AF_T_ removes only the template structures. AF_S_ removes only homologous sequences. AF_ST_ removes all homologous templates and sequences beyond a sequence identity threshold. Each result is the average of three independent simulations. We also provide the MSA depth which is used when removing sequences in AFS and AFST.

| System<br>(Ref.<br>PDB) | Identity<br>threshold<br>(%) | $AF_T$ | | $AF_S$ | | $AF_{ST}$ | | MSA<br>Depth |
| --- | --- | --- | --- | --- | --- | --- | --- | --- |
|  |  | Global | BS | Global | BS | Global | BS |  |
| Thrombin<br>(2ZFF) | 100 | 2.56 | 2.40 | 2.52 | 2.40 | 2.74 | 2.56 | 16669 |
|  | 70 | 1.38 | 0.45 | 2.65 | 2.44 | 1.36 | 0.34 | 16452 |
|  | 30 | 1.39 | 0.43 | 2.65 | 2.47 | 2.75 | 0.35 | 14169 |
|  | 20 | 1.37 | 0.42 | 2.81 | 2.53 | 2.81 | 1.85 | 12613 |
|  | 10 | 1.39 | 0.45 | 2.84 | 2.58 | 7.71 | 1.69 | 3726 |
|  | 5 | 1.57 | 0.45 | 2.73 | 2.61 | 31.71 | 16.29 | 89 |
| CDK2<br>(1H1Q) | 100 | 3.58 | 2.64 | 3.65 | 2.64 | 3.71 | 2.61 | 11468 |
|  | 70 | 3.58 | 2.59 | 3.84 | 2.65 | 3.65 | 2.68 | 10844 |
|  | 30 | 3.66 | 0.80 | 2.27 | 2.42 | 4.05 | 2.61 | 9141 |
|  | 20 | 3.62 | 0.64 | 2.38 | 0.70 | 2.35 | 0.82 | 5190 |
|  | 10 | 3.66 | 0.64 | 3.62 | 0.57 | 22.96 | 9.70 | 38 |
|  | 5 | 4.46 | 2.77 | 4.24 | 2.54 | 20.84 | 15.05 | 1 |
| PTP1B | 100 | 0.37 | 0.92 | 0.36 | 0.92 | 0.38 | 0.94 | 4467 |
| (2QBS) | 70 | 1.09 | 0.97 | 0.36 | 0.92 | 1.36 | 0.99 | 3793 |
|  | 30 | 0.76 | 0.38 | 0.40 | 1.03 | 0.97 | 0.91 | 6897 |
|  | 20 | 1.11 | 0.34 | 0.51 | 1.14 | 1.44 | 1.02 | 3028 |
|  | 10 | 0.89 | 0.29 | 0.60 | 1.50 | 2.76 | 1.94 | 222 |
|  | 5 | 1.01 | 0.91 | 0.63 | 1.62 | 17.14 | 14.03 | 3 |

### AF2 customization

AF2 employs both structural templates as well as Multiple Sequence Alignments (MSA) in order to predict structures. In our custom version of AF2 we systematically removed all template structures and sequences above 30% sequence identity from the database used to build these models. This ensured that our benchmark reflects a prospective application of AF2 in a drug discovery project: namely a situation where no high-quality homology model building would be possible due to the lack of availability of a high-sequence identity template.

To perform this realistic benchmark, we performed extensive customization of the AF2 code, as described in detail in Methods. Briefly, the new version is now capable of removing either structural templates, or sequences, or both, from the AF2 database, based on a user-defined sequence identity threshold (see Figure 1). Models are then created taking into consideration only structures and sequences below this identity threshold.

**Figure 1.**
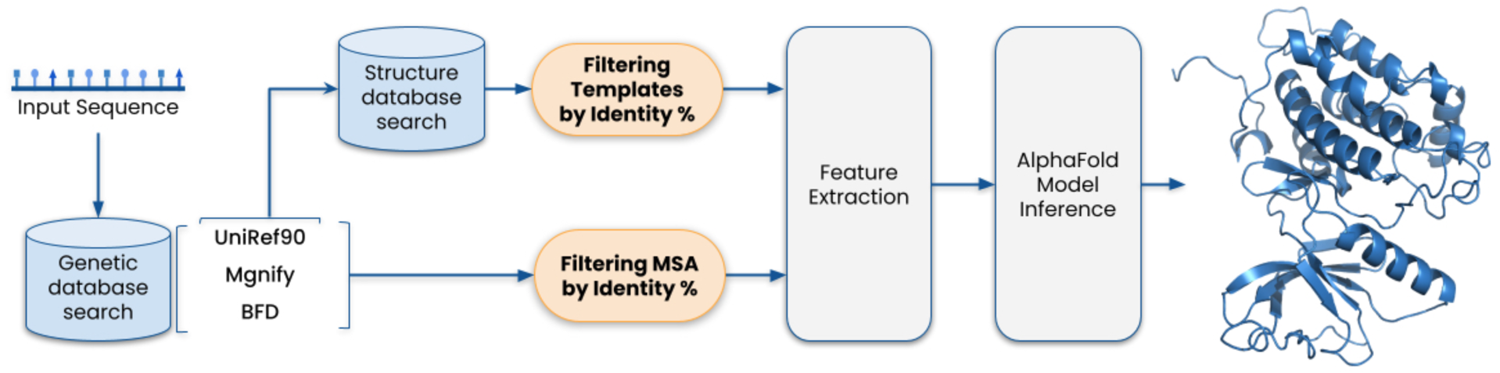
Overview of the customized AF2 pipeline. Sections framed in orange are customizations with respect to the original AF2 workflow. The model uses evolutionary related protein sequences and amino acid residue pairs (Feature Extraction) to iteratively pass the information to an end-to-end transformer-based neural network (AlphaFold Model Inference), in order to generate a 3D structure.

Table 1 shows three examples where we analyzed the effect on model accuracy of systematically removing sequences (AF_S_) or templates (AF_T_) or both (AF_ST_) above different identity thresholds from the AF2 database. Thus, in the first column, AF_ST_ reflects the experiment where both sequences and templates were removed above the identity threshold. AFS reflects the scenario where only sequences below a given identity threshold are included in the MSA. In the AFT column, the MSA is not filtered, while the template structures are culled based on identity. Interestingly, we see some significant fluctuations which, a priori, might seem counterintuitive.

For example in Thrombin, when going from high to low identity filters, we observe an improvement in prediction accuracy when removing structural templates of high sequence identity. The reason for this effect is that these high identity templates (e.g. 6C2W) present an ordered alpha helical structure in the active site (see Figure S1), which corresponds to a disordered loop in 2ZFF (the structure used in the original FEP+ study^11^ and used as a reference for RMSD calculations here).

The fine tuning between templates and MSA network weights is the result of extensive deep learning training, and not easy to rationalize. A case in point is the low binding site RMSDs in CDK2 when removing structural templates or sequences above 30%, compared to the higher RMSD when high sequence identity templates are used. In this regard AF2 has the impressive ability to create low RMSD models based solely on sequence evolutionary data, even when removing basically all structural templates. Similarly, when reducing the depth of the MSA (second column, 5% identity threshold), AF2 is able to produce models with low RMSD values when depending exclusively on structural templates. This scenario would be equivalent to developing a model using state-of-the-art homology modeling algorithms. In our three examples, i t is only when both sequences and structures are culled from the database beyond 5% sequence identity (or 10% in some cases) that it becomes impossible to produce high-quality models. Overall, however, the performance of the algorithm when using only limited data is remarkable. AF2 is an extremely robust predictor, capable of extracting structural information from low-identity templates and sequences to create highly accurate models.

### Structural Model Accuracy

For developing models for the FEP+ benchmark, we chose an identity threshold of 30% to remove sequences and templates from the database. The resulting accuracy of all the models generated with this custom AF2 implementation (henceforth named AF2_30_) is described in Table 2. We captured the accuracy of the model using a variety of metrics, including: 1) global RMSD of the model with respect to the crystal structure used in the original FEP benchmarks; 2) bindingsite RMSD with respect to the crystal structure; and 3) RMSD of the ligand modeled into the apo AF2_30_ structures (see Methods for description on how ligand poses in AF2_30_ structures were determined).

**Table 2.** RMSD values of models produced with AF2_30_ models, aligned with the reference crystal structure. Global and binding site RMSD values were calculated with Cα-atoms only. The binding site includes all residues within 5Å from the ligand in the reference structure. Ligand RMSD values were calculated for all heavy atoms, following an alignment of the binding site residues. The PLDDT score reflects a confidence measurement in the accuracy of the structure, as reported by the AF2 algorithm.

| Target | Reference structure | Global RMSD | Binding Site RMSD | Ligand RMSD | AF2<br>confidence<br>score (PLDDT) |
| --- | --- | --- | --- | --- | --- |
| A2A | 4EIY | 2.09 | 0.56 | 2.18 (0.46*) | 86.91 |
| B1AR | 3ZPQ | 1.01 | 0.56 | 0.84 | 73.86 |
| BACE | 4DJW | 1.92 | 1.20 | 1.14 | 83.67 |
| CDK2 | 1H1Q | 3.66 | 0.80 | 0.61 | 88.45 |
| CDK9 | 4BCI | 1.64 | 1.33 | 1.16 | 86.61 |
| HSP90 | 3FT8 | 0.88 | 1.24 | 1.48 | 83.32 |
| ERK2 | 5K4I | 1.47 | 1.22 | 0.71 | 90.03 |
| JAK2 | 3E64 | 0.98 | 1.20 | 0.66 | 86.88 |
| JNK1 | 2GMX | 1.40 | 0.74 | 0.90 | 81.46 |
| MCL1 | 4HW3 | 1.12 | 1.20 | 1.14 | 65.07 |
| P38 | 3FLY | 2.85 | 1.54 | 0.88 | 88.97 |
| PTP1B | 2QBS | 0.76 | 0.38 | 0.85 | 79.96 |
| Thrombin | 2ZFF | 1.53 | 0.44 | 0.35 | 83.86 |
| Tyk2 | 4GIH | 1.14 | 0.65 | 0.46 | 81.93 |

Overall the accuracy of the AF2_30_ structures is excellent. All global RMSD values (calculated using all residues visible in the original crystal structures) are below 2.85Å, and 75% of the models have RMSD values below 2Å. Given the dependence of FEP calculations on accurate description of the protein-ligand interactions, a more relevant RMSD value for our benchmarking purposes is binding site RMSD. Here too the values are excellent across the board, with all values below 1.54Å and 50% of all models below 1Å. In addition, all except for one model are classified as highly reliable, as evidenced by confidence scores (PLDDT) that range between 70 and 90. Together, these results suggested to us that FEP+ calculations using these models would have a high likelihood of success.

Moreover, we measured the RMSD of the ligand in our AF2_30_ structure compared to the crystal structure pose. The superimposed complexes are shown in Figure 2. With one exception, all RMSD values are below 1.4Å. The one exception is the A2A receptor, which has an RMSD of 2.18Å. However, this includes a flexible part of the molecule that rearranges in response to a clash with a side-chain residue. If only the rigid core of the molecule is considered the RMSD drops to 0.46Å. Similar observations regarding the prediction of this ligand have been made previously^19^.

**Figure 2.**
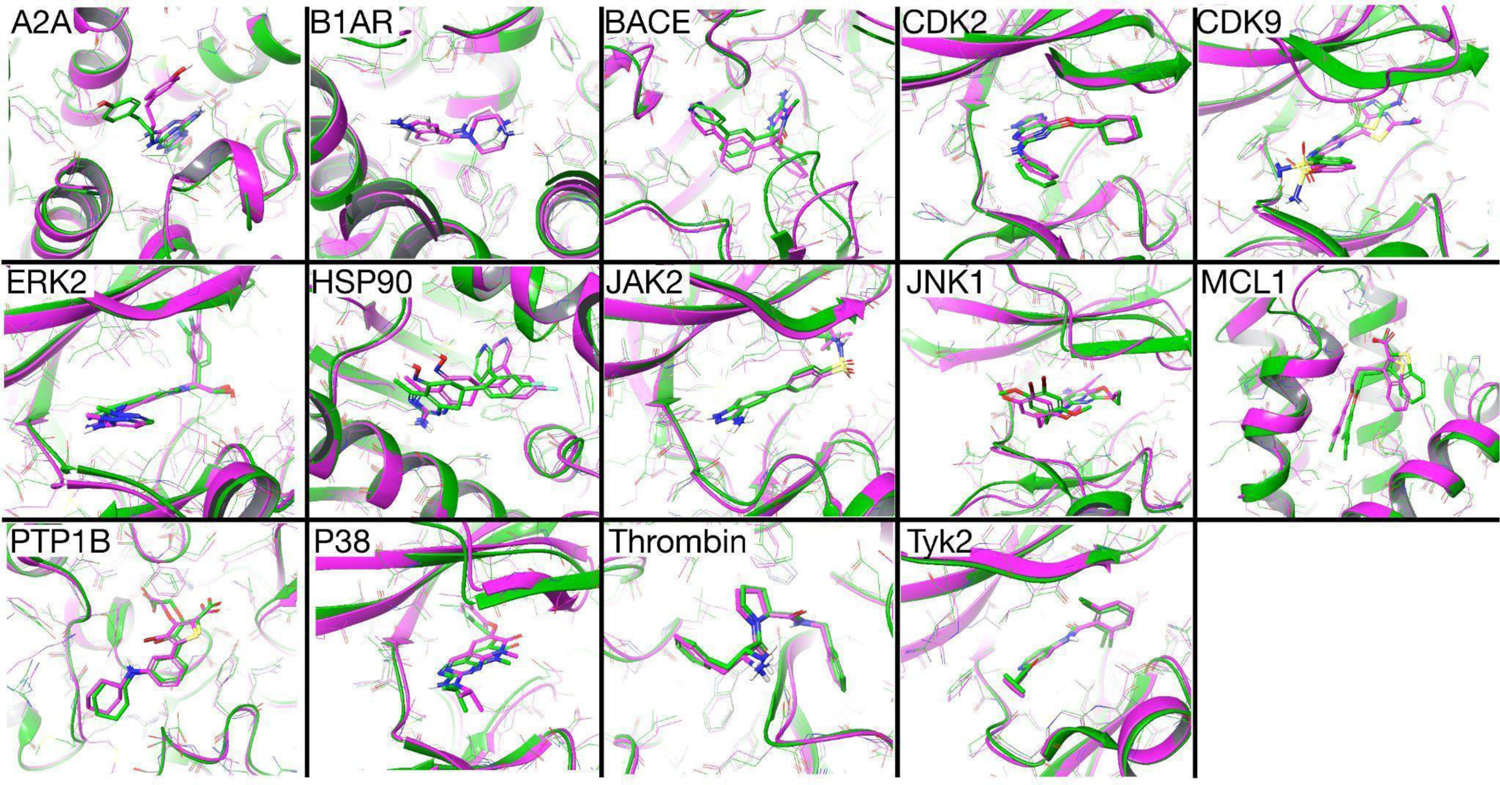
Superposition of AF2_30_ models (green) on corresponding crystal structures (magenta). Refer to Table 2 for a description of the accuracy of these models compared to the crystal structures.

We also attempted a comparison of the accuracy of AF2_30_ models to those produced with current state-of-the-art homology modeling methods using low sequence identity templates (>30%). In all cases a single template approach was used, selecting the template with the highest identity used by AF2_30_ (within a maximum identity of 30%, see Table S1 for an overview of sequence and template IDs used in the homology modeling exercise). Table 3 shows the global and binding site RMSD values obtained by three different (and widely used) homology modeling methods: Prime^20,21^, iTasser^22^ and SwissModel^23^. The superpositions of the resulting models on the reference crystal structures are shown in supplementary figure S2. In the vast majority of cases AF2_30_ models were superior to those created with the homology modeling methods.

**Table 3.** Comparison of RMSD values of all models - obtained with three different homology modeling programs and AF2_30_, aligned with the reference crystal structure. Global all-atom RMSD (All), Cα carbon RMSD (CA) and Cα binding site RMSD values (BS) are reported in Å.

| System | Prime |  |  | I-Tasser |  |  | SwissModel |  |  | AF2<sub>30</sub> |  |  |
| --- | --- | --- | --- | --- | --- | --- | --- | --- | --- | --- | --- | --- |
|  | All | CA | BS | All | CA | BS | All | CA | BS | All | CA | BS |
| A2A | 10.42 | 10.09 | 10.5 | 2.76 | 2.22 | 0.40 | 14.15 | 13.60 | 3.65 | 2.34 | 2.09 | 0.56 |
| B1AR | 4.30 | 3.51 | 2.05 | 4.07 | 3.36 | 1.88 | 4.34 | 3.81 | 2.06 | 1.39 | 1.01 | 0.56 |
| BACE | 6.65 | 6.04 | 2.21 | 9.64 | 9.12 | 2.61 | 5.65 | 4.94 | 2.61 | 2.23 | 1.90 | 1.20 |
| CDK2 | 7.15 | 6.53 | 2.51 | 6.12 | 5.81 | 1.65 | 7.68 | 7.35 | 1.87 | 4.08 | 3.66 | 0.80 |
| CDK9 | 6.35 | 5.47 | 1.20 | 2.26 | 1.59 | 1.32 | 5.63 | 4.88 | 1.19 | 2.18 | 1.64 | 1.33 |
| ERK2 | 12.99 | 12.59 | 2.47 | 15.32 | 14.85 | 1.09 | 7.92 | 7.40 | 1.22 | 1.89 | 1.47 | 1.22 |
| HSP90 | 20.77 | 20.15 | 14.1 | 9.13 | 9.04 | 1.38 | 8.04 | 7.61 | 2.83 | 1.32 | 0.88 | 1.24 |
| JAK2 | 7.10 | 7.07 | 1.06 | 13.42 | 13.56 | 20.8 | 5.46 | 5.01 | 6.50 | 1.60 | 0.98 | 1.20 |
| JNK1 | 13.25 | 12.61 | 0.95 | 11.30 | 10.71 | 0.95 | 8.23 | 7.79 | 0.94 | 2.04 | 1.40 | 0.74 |
| MCL1 | 3.99 | 3.03 | 2.27 | 4.45 | 4.19 | 1.55 | 4.63 | 4.45 | 1.68 | 1.67 | 1.12 | 1.20 |
| P38 | 13.90 | 13.44 | 5.41 | 11.66 | 11.27 | 1.28 | 7.35 | 6.39 | 2.42 | 3.22 | 2.85 | 1.54 |
| PTP1B | 8.01 | 7.41 | 1.60 | 5.25 | 4.56 | 1.78 | 2.80 | 2.00 | 1.53 | 1.37 | 0.75 | 0.38 |
| Thrombin | 7.23 | 6.59 | 4.26 | 5.56 | 4.84 | 3.02 | 4.67 | 3.79 | 4.35 | 1.63 | 1.53 | 0.44 |
| Tyk2 | 7.00 | 6.45 | 0.86 | 3.49 | 2.85 | 1.18 | 5.38 | 4.90 | 0.83 | 1.66 | 1.13 | 0.65 |

It is important to point out that we did not include an explicit docking step in the preparation of the input for the FEP calculations. Rather, we simply translated the ligand coordinates from the crystal structure into the model based on superposition, followed by a brief optimization step using Prime^24^.

### FEP results

The original FEP+ benchmark study used between 11 and 42 ligands per target, resulting in a total number of perturbations between 16 and 71^11^. Due to limited computational capacity, we did not attempt to reproduce the entire dataset obtained in the original work. Rather, for each of the targets in the original FEP+ benchmark studies, we selected a representative subset of perturbations to reproduce using the AF2_30_ structures (between 7 and 18). In order to ensure a fair comparison, we made sure that within our chosen subset the mean unsigned error (MUE) of the predicted ΔΔG between pairs of compounds was similar to the MUE of the entire dataset. In other words, the selected perturbations included those that were predicted with very high accuracy using FEP+, as well as perturbations for which significant errors were reported. In addition, we calculated correlations between experimental and calculated ΔG values by calculating additional edges in the subset maps, in order to obtain cycle-closure corrected results.

The aggregated results of the FEP+ benchmark are reported in Table 4. Detailed results are available in Supplementary table S2 to S5. In general, the accuracy of FEP+ calculations performed using AF230 models, in terms of MUE, is not statistically different from the error obtained with crystal structures. The average error across all targets using AF2_30_ models is 1.04 kcal/mol, compared to 1.01 kcal/mol for crystal structures. The same general trend holds true for targets reported in the original FEP paper (MUE of 0.90 kcal/mol for AF2_30_ models compared to 0.97 kcal/mol for crystal structures), for fragment datasets (MUE of 0.93 kcal/mol for AF2_30_ models compared to 1.35 kcal/mol for crystal structures), and for GPCRs (MUE of 1.22 kcal/mol for AF2_30_ models compared to 0.90 kcal/mol for crystal structures). In the case of selectivity studies instead of MUEs compound affinities were reported, making a direct comparison difficult. However, the MUE values calculated for the subset of compounds tested (1.32 kcal/mol) here are similar to the average values reported for the full dataset (1.05 kcal/mol), suggesting that here too AF2_30_ models perform similarly to crystal structures. Since the identity of the maps calculated here is different from those reported in the original benchmarks, a direct comparison on R^2^ values is not possible. However, in 11 out of 16 cases the observed R^2^ falls within the range of expected R^2^ values for FEP-predicted binding affinities and experimental results with assumed RMSEs of 1.1 kcal/mol, demonstrating the utility of these results for rank ordering compounds based on predicted affinities.

**Table 4.**
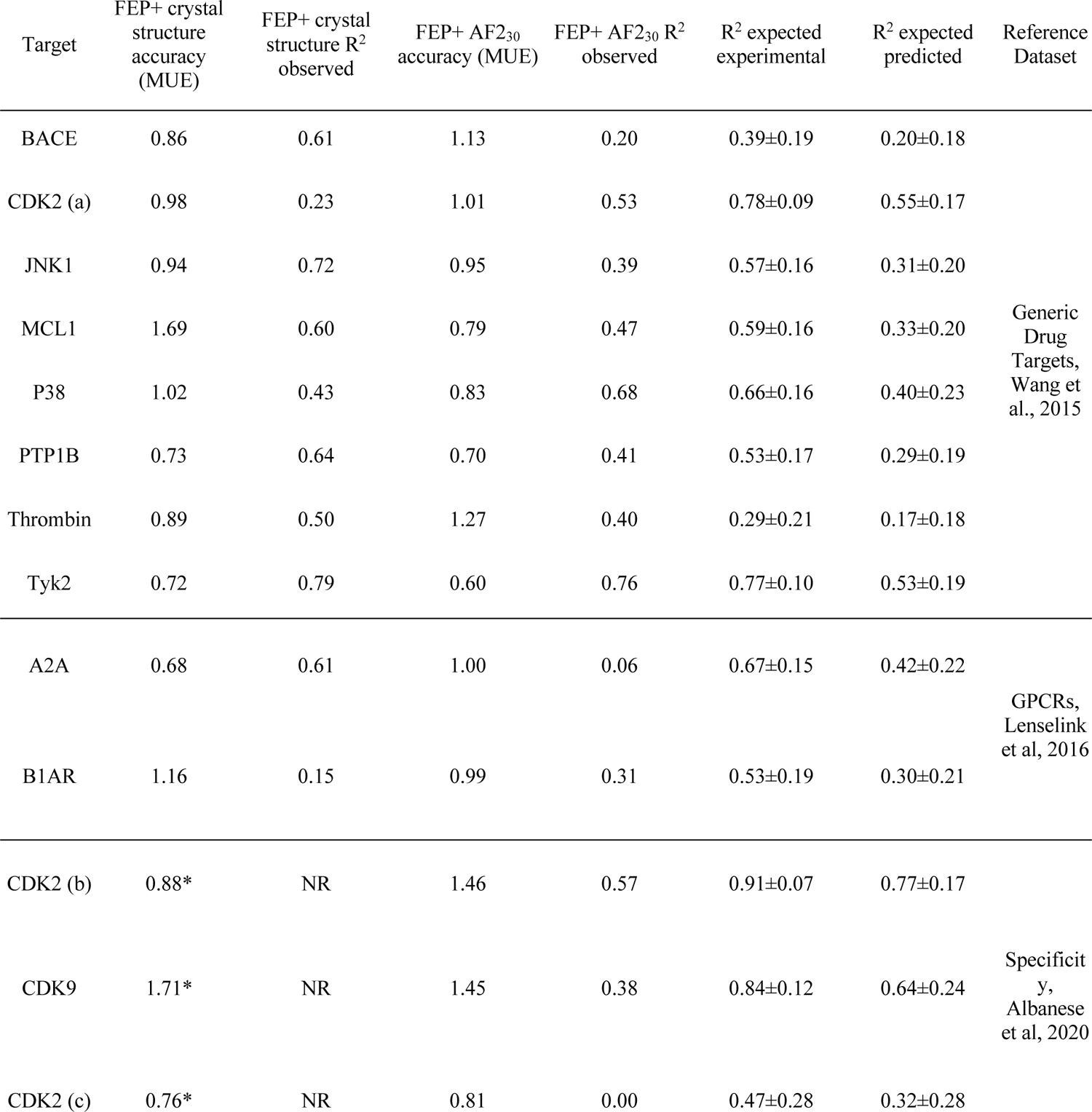
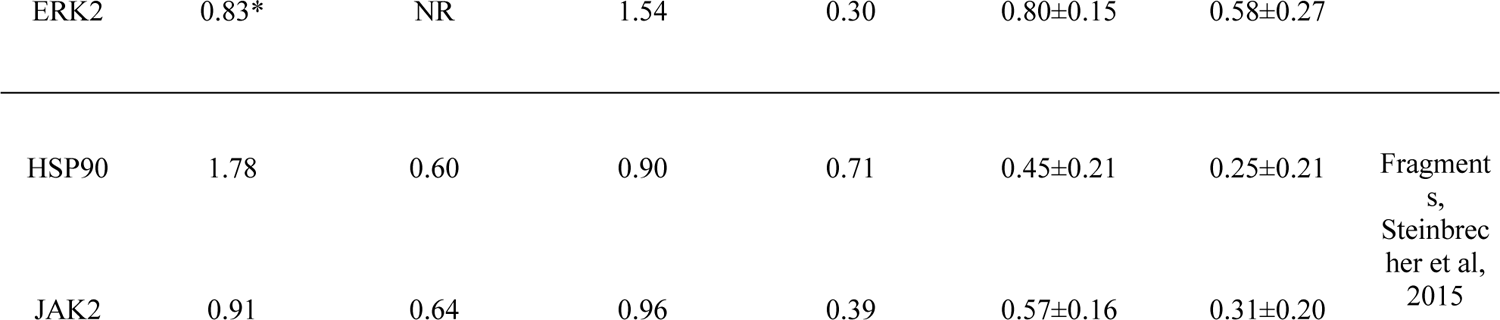
Summary of FEP results. All MUE values (in kcal/mol) were calculated from the individual perturbations as reported in the supplementary information of the Wang^11^ and Lenselink^15^ studies. Results for individual perturbations in the Steinbrecher^14^ study were provided by the authors upon request. MUE values for the specificity set (indicated with an asterix) were only available for the entire dataset in the original FEP+ benchmark. R^2^ values for results obtained here were calculated using the subset map (column 4), while R^2^ values for results obtained with crystal structures were calculated using maps of the entire dataset. Expected R^2^ values between FEP-predicted binding affinities and experimental results (R^2^ expected predicted) and expected correlation coefficient between two experimental measurements of binding affinities (R^2^ expected experimental), with assumed RMSEs of 1.1 and 0.4 kcal/mol for FEP-predicted binding affinities and experimental data, respectively, are also shown^11^. The three CDK2 datasets involve different chemical series, one (a) obtained from Wang et al ^11^ and two (b and c) from Albanese et al.^16^

Finally, Figure 3 shows the correlation of predicted ΔG against experimental ΔG for all 16 series. Most compounds are predicted within 1 kcal/mol of their experimental affinity (102 out of 138), and all but three compounds are predicted within 2 kcal/mol of the experimental affinity.

**Figure 3.**
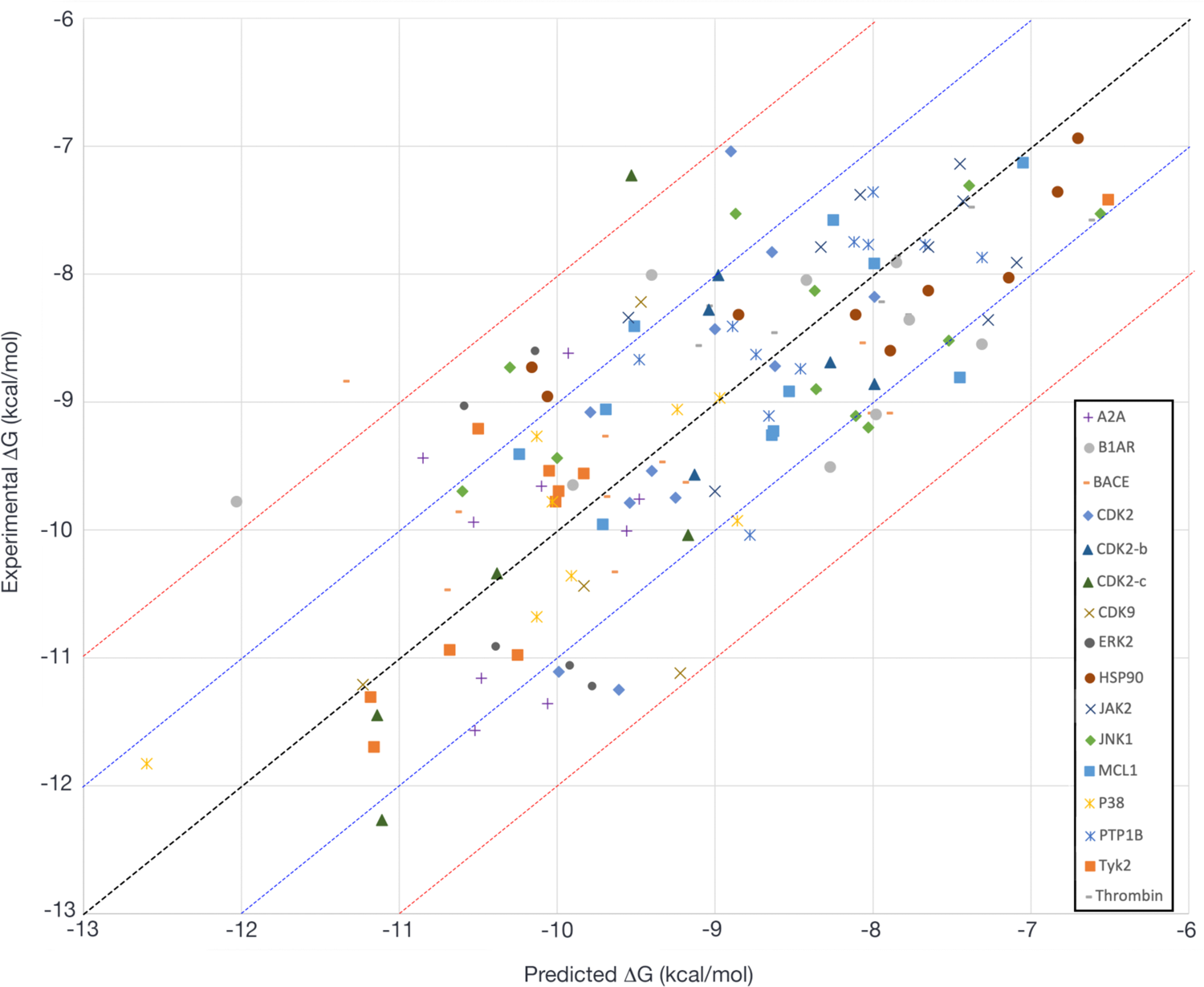
Predicted ΔG obtained with FEP+ plotted against experimental ΔG, for all 16 compound series studied here. Most of the predicted values for the 138 ligands fall within <1.0 kcal/mol of the experimental results (blue diagonal lines) and all but three compounds are predicted within 2.0 kcal/mol of the predicted affinity (red diagonal lines).

## Discussion

There is an urgent interest in determining the potential of deep learning protein structure predictions techniques in solving practical problems in chemistry and biology. One obvious domain of applicability is the discovery and optimization of novel small molecule therapeutics, where the availability of accurate target structures continues to seriously hinder drug discovery projects. AF2 (and other related techniques) could drastically impact this situation. To address this, we have tested how AF2-predicted models could substitute crystal structures using a gold standard affinity prediction method, FEP+. To this end, we designed a study where we reproduced a representative group of calculations from the original FEP+ benchmark studies^11,14,15,16^. Moreover, in order to mimic a realistic prospective scenario, we limited AF data sources (both structural templates and sequences) to an identity threshold of 30%, a value previously considered to be the ‘twilight’ zone of homology modeling accuracy^25^. This identity threshold has consistently been shown to provide mid-to low-quality structures in blind CASP competitions^26^.

The quality of the AF2_30_ predictions of all targets included can be considered high, as reflected by the high value of the confidence scores (see reported PLDDT values in Table 2). Indeed, global and binding site RMSD values are within the range observed when comparing different structures for the same target.

We systematically assessed the effects of imposing a sequence identity threshold on the data fed into the AF2 algorithm, in three different targets. As expected, in the control situation where all structural templates and sequences similar to the target of interest are removed (using a 5% identity threshold), only low-quality models are produced. This indicates that the recognition of structures is not deeply coded within the deep learning model, a concern since all three structures were used for the training of the model. In more realistic scenarios, i.e., the presence in the database of structural templates with up to 30% identity, and/or a large number of sequences available for the

MSA, AF2 performs extraordinarily well. Still, accuracy values can fluctuate in manners that are quite unpredictable in response to removing parts of the data from the algorithm. This effect has already been pointed out by the AF2 authors in their original manuscript, where they advise not to remove any data source from consideration^3^. Indeed, a detailed explanation of the effects of changes in the structural templates and/or sequence database on model quality remains difficult, and a more comprehensive benchmark is warranted. We would recommend, however, that as an estimator of the model quality, it is useful to assess the RMSD among the top structural templates selected by AF2. In cases where this RMSD is high the availability of a large MSA might be preferred to that of high-identity structural templates. In this sense, we observed that the potential of using MSA data alone is in most cases enough to exceed the accuracy levels of models produced by other homology modeling software. While all data produced here was obtained without any user intervention, and as such it is likely that model quality can be improved through customization of sequence alignment and model building parameters, there is clearly a large gap between AF2 and the methods compared to here (as seen already in the recent CASP competitions^27^).

While further studies regarding the ability of AF2 to produce high quality models in cases where no good structural templates are available are needed, the main goal of this study was to assess the utility of such models to support computationally driven lead-optimization, for example using FEP. The results obtained here comprehensively show that - in the limit of the ability to predict ligand binding modes given an apo structure - AF2_30_ derived models are just as reliable as crystal structures in terms of predictive accuracy. The MUE of the individual perturbations for calculations done with AF230 are comparable with those done with crystal structures, and in many cases the R^2^ values obtained for cycle-closed maps exceed or are similar to the expected values for well-behaving FEP calculations. This suggests that the application of FEP in prospective discovery projects will depend less on the availability of experimentally derived structures, and more on the intrinsic limitations of the method, including the accuracy of the forcefield, the ability to sample relevant conformational states, and the accurate treatment of different charged states^28^.

Given the fact that we omitted a docking stage into our computational workflow, and relied on superposition to generate protein-ligand complexes, the results obtained here present an upper limit of what can be achieved in terms of FEP calculations, reflecting situations where the ligand pose can be predicted with high accuracy. In many cases (e.g., kinases) specific recognition motifs can be identified in the ligand (e.g., the hinge binding part of the compound) and using these criteria the ligand can often be placed into the binding site unambiguously. In other cases, uncertainty in the docking step has the potential to significantly affect any downstream FEP calculations. Here, next generation docking tools for the purpose of pose predictions can ensure reliable starting conformations for FEP calculations^29,30^. In addition, FEP calculations using multiple starting structures can be used to validate docking studies. However, while an in-depth assessment of the utility of AF230 structures for docking calculations is warranted, it is beyond the scope of this current work.

It is possible that the high accuracy of the current calculations is partially due to improvements to the FEP+ algorithm since the initial publication of the method in 2015, including the use of a different ensemble (μVT as opposed to NPT). Indeed, it is likely that the improved quality of the currently used forcefield^31^ and ensemble would lead to a small but significant improvement in calculations using crystal structures as well. To test this hypothesis, we carried out FEP calculations with the current version of the software, for two targets where the AF2_30_ models outperformed the original crystal structure-based calculations by the largest degree (targets MCL1 and P38). Indeed, with the latest version of the method the MUE was significantly reduced (MCL1 from 1.69 kcal/mol to 1.05 kcal/mol and P38 from 1.02 kcal/mol to 0.67 kcal/mol). Results are presented in Supplementary Table S6. Despite this effect of the improved algorithm, it can be concluded that in general the quality of the results of using deep learning derived models is very close to what can be expected to be obtained using crystal structures, using the currently available implementation of FEP+. These advances have the potential to dramatically increase the domain of application for FEP in prospective drug discovery settings.

## Methods

### Dataset

A total of 14 targets were selected for use in this FEP benchmark. This included eight pharmaceutically relevant targets studied in the original FEP+ benchmark^11^, two target sets previously studied for specificity prediction^16^, two targets previously studied in a fragment benchmark^14^, and two targets previously studied in a GPCR-specific benchmark^15^. PDBs used to model the targets in those studies are shown in Table S1.

### AlphaFold2 customization

AF2 makes use of a data pipeline in its prediction algorithm. During the first stage, the algorithm identifies homologues of a query sequence and makes use of Hidden Markov Models methods to construct a multiple sequence alignment (MSA). This MSA is used to derive evolutionary information of the target protein. To generate the MSA the current protocol searches multiple databases: BFD^32,33^, Mgnify^34^ and Uniref90^35^. MSAs outputs are de-duplicated and combined and used to feed the neural network. In a second stage, the Uniref90 multiple sequence alignment is used to find structural templates to feed into the neural network. The top 20 of these templates are selected and then AF2 picks the top four, sorted by the expected number of correctly aligned residues (the “sum_probs” feature generated by HHSearch).

At present, structural templates (from the PDB70 database) can be filtered out only by PDB release date. This limitation makes it difficult to anticipate how AF2 would behave in those cases where very little data is available (i.e., no high-identity homolog templates and sequences for a protein of interest). To overcome these restrictions, we modified the AF2 code to filter data by identity threshold but without altering its original workflow, enabling the user to restrict the data that AF2 can use from each database. Data restriction was accomplished by modification of the multiple sequence alignments features. Before stacking the genetic search outputs, we computed the identity of each hit and excluded those with a higher identity than the desired threshold. Moreover, we applied this strategy at different levels: i) by removing both templates and restricting the MSA by filtering all databases; ii) by only reducing the size of the MSA by filtering BFD, MGnify, and Uniref90 databases; iii) by only removing templates through filtering PDB70.

### Modeling of reference ligand

The ligand was introduced into the apo AF2_30_ model by aligning the model with the crystal structure used for original FEP calculations, and through introduction of the aligned ligand pose through superposition. In some cases this resulted in significant clashes between the ligand and the protein. Optimization of the resulting protein ligand complex using Maestro’s Refine Protein Ligand complex utility^24^ (run using default settings) was able to resolve these clashes, but in some cases resulted in significant deviation of the ligand with respect to the crystal structure (see Table 2).

### Homology modeling

We used three widely employed homology modeling methods to compare with AF230: Prime^20,21^, I-Taser^22^ and Swiss Model Server^23^. In each case provided only a single structural template along with the target sequence. For each system, we inspected the templates used in AF2_30_ and chose the one with the highest identity (within the identity threshold of 30%). If more than one template had a maximum identity percentage, the one with higher sum_probs was chosen since this is the metric used by AF2 to rank sequences in the MSA. Templates used for each model are shown in the supporting Information. For the sake of reproducibility, all programs were run using default parameters.

Prime^20,21^ is a suite of programs for protein structure prediction. The Homology Modeling workflow consists of template identification, alignment, and model building. ClustalW was used to perform the alignment of the template and input sequence. The loops of the models were built using knowledge-based methods.

I-TASSER (Iterative Threading ASSEmbly Refinement) is a hierarchical approach to protein structure prediction and structure-based function annotation^22^. The I-TASSER Server was ranked as the No 1 server for protein structure prediction in recent community-wide CASP experiments (https://predictioncenter.org). To build the homology models both the fasta sequence extracted from the UniProt database and a template PDB ID was provided. We selected the model with higher C-score among the 5 models to compute RMSD against the crystal.

Swiss Model^23^ is a fully automated and widely used protein structure homology-modeling server. To generate the homology model, we provided the UniProt ID of the target protein and the AF2 best template PDB (removing HETATMS and chains different to the ones specified in Table S1).

#### Ligand datasets

For each of these targets the ligands used in the FEP+ benchmarks were downloaded from Chembl^36^, or when not available sketched by hand using Maestro^37^. Affinity values were obtained from Supporting Information in the original FEP+ publications. Ligands were introduced into the binding site by flexible alignment with the optimized reference ligand in the AF2_30_ model, using Maestro’s ligand alignment function. In some cases it was necessary to manually rotate aromatic rings and reposition R-groups to avoid steric overlap with the protein.

Perturbation selection: For each target ten perturbations were selected for evaluation with FEP+. We carefully chose these to include perturbations with low and high error in the original benchmarks. For example, the MUE of the perturbations selected (as previously reported by Wang et al. 2015^11^) was 0.97 kcal/mol, compared to 0.92 kcal/mol for the entire dataset. The results of all calculated perturbations are provided as Supplementary Information.

FEP+ calculations: Calculations were run using the 2021-4 version of FEP+^11^. Default settings were used, which includes 12 lambda windows, 5ns sampling time per lambda with replica exchange, the μVT ensemble, and use of the OPLS4 forcefield^31^. Force field parameters were calculated where required using the Force Field builder tool in Maestro^37^.

## Author Contributions

The manuscript was written through contributions of all authors. All authors have given approval to the final version of the manuscript.

## Funding Sources

This work was supported by Grant PTQ2018-009991 funded by MCIN/AEI/ 10.13039/501100011033, and additional funding from RA Capital.

## ABBREVIATIONS

FEP: Free Energy Perturbation
AF2: AlphaFold2
MSA: Multiple Sequence Alignment
A2A: Adenosine receptor 2A
B1AR: beta1 adrenergic receptor
BACE: Beta-secretase
CDK2: Cyclin-dependent kinase 2
CDK9: Cyclin-dependent kinase 9
HSP90: heat shock protein 90
ERK2: extracellular signal-regulated kinase 2
JAK2: Janus kinase 2
JNK1: c-Jun N-terminal kinase 1
MCL1: Induced myeloid leukemia cell differentiation protein
ND: Not Determined
P38: p38 mitogen-activated protein kinase
PTP1B: protein-tyrosine phosphatase 1B
RMSD: root mean square deviation
MUE: mean unsigned error
RMSE: root mean square error;

## CODE AVAILABILITY

Our custom AlphaFold implementation to remove sequences and/or templates has been published on GitHub under Apache License 2.0: https://github.com/hemahecodes/alphafold.

## Notes

### Competing Interest Statement

The authors have declared no competing interest.

### Summary of Updates

No changes to the manuscript, but corrected the title in Manuscript Basics and changed a typo in an authors name

