## Supplementary Information for "Are Deep Learning Structural Models Sufficiently Accurate for Free Energy Calculations? Application of FEP+ to AlphaFold2 Predicted Structures"

Supplementary Tables

| System | Uniprot ID | Template PDB ID |
|---|---|---|
| A2A | P29274 | 3V2Y:A |
| B1AR | P08588 | 6OIJ:R |
| BACE | P56817 | 5N7Q:A |
| CDK2 | P24941 | 3KK8:A |
| CDK9 | P50750 | 4QNY: |
| ERK2 | P28482 | 2OWB:A |
| HSP90 | P07900 | 6RKW:B |
| JAK2 | O60674 | 4XEY:B |
| JNK1 | P45983 | 5G6V:A |
| MCL1 | Q07820 | 6FBX:A |
| p38 | Q16539 | 5EFQ:C |
| PTP1B | P18031 | 2NLK:A |
| Thrombin | P00734 | 2XXL:A |
| Tyk2 | P29597 | 2J0K:B |

Table S1. Target Uniprot and template PDB IDs used for creating homology models.

| | Code ligand A | Code ligand B | Exp ΔΔG | Crystal Structure ΔΔG | Crystal Structure Absolute Error | $AF2_{30}$ ΔΔG | $AF2_{30}$ Absolute Error |
|---|---|---|---|---|---|---|---|
| BACE | CAT-13h | CAT-17i | 0.16 | 0.14 | 0.02 | -0.03 | 0.19 |
| | CAT-13d | CAT-13h | 0.84 | 1.46 | 0.62 | 1.72 | 0.88 |
| | CAT-13a | CAT-17i | -0.63 | -0.76 | 0.13 | 2.69 | 3.32 |
| | CAT-13o | CAT-17i | -0.93 | -1.08 | 0.15 | -1.7 | 0.77 |
| | CAT-13o | CAT-17h | -1.79 | -1.98 | 0.19 | -1.52 | 0.27 |
| | CAT-13g | CAT-17i | -0.38 | 0.84 | 1.22 | -1.5 | 1.12 |
| | CAT-13d | CAT-13f | 1.38 | 0.13 | 1.25 | 2.7 | 1.32 |
| | CAT-13g | CAT-17g | -0.65 | 0.86 | 1.51 | -2.72 | 2.07 |
| | CAT-17g | CAT-17c | -0.12 | -1.86 | 1.74 | -0.99 | 0.87 |
| | CAT-13d | CAT-13i | 1.2 | -0.59 | 1.79 | 0.72 | 0.48 |
| | | | | | MUE 0.86 | | MUE 1.13 |

| | Code ligand A | Code ligand B | Exp ΔΔG | Crystal Structure ΔΔG | Crystal Structure Absolute Error | $AF2_{30}$ ΔΔG | $AF2_{30}$ Absolute Error |
|---|---|---|---|---|---|---|---|
| CDK2 | 20 | 1h1q | 0.539 | 0.61 | 0.071 | 0.64 | 0.101 |
| | 31 | 32 | -0.211 | 0.01 | 0.221 | 0.13 | 0.341 |
| | 1oiy | 31 | 0.246 | 0.01 | 0.236 | 0.38 | 0.134 |
| | 28 | 31 | 1.573 | 1.3 | 0.273 | 0.28 | 1.293 |
| | 1oiy | 1h1q | 1.606 | 0.54 | 1.066 | 1.47 | 0.136 |
| | 28 | 26 | 2.681 | 1.48 | 1.201 | 1.28 | 1.401 |
| | 17 | 21 | -0.787 | 0.57 | 1.357 | 0.62 | 1.407 |
| | 1h1s | 26 | 2.815 | 1.2 | 1.615 | 0.56 | 2.255 |
| | 1oiu | 1h1q | 0.904 | 2.75 | 1.846 | 2.05 | 1.146 |
| | 17 | 1h1q | -1.138 | 0.78 | 1.918 | 0.71 | 1.848 |
| | | | | | MUE 0.98 | | MUE 1.01 |

| | Code ligand A | Code ligand B | Exp ΔΔG | Crystal Structure ΔΔG | Crystal Structure Absolute Error | $AF2_{30}$ ΔΔG | $AF2_{30}$ Absolute Error |
|---|---|---|---|---|---|---|---|
| JNK1 | 18632-1 | 18624-1 | 0.59 | 0.6 | 0.01 | 0.65 | 0.06 |
| | 18636-1 | 18624-1 | -0.98 | -1.07 | 0.09 | 1.15 | 2.13 |
| | 18633-1 | 18624-1 | 0.68 | 0.78 | 0.1 | 0.8 | 0.12 |
| | 18636-1 | 18625-1 | -0.59 | -0.3 | 0.29 | 0.7 | 1.29 |
| | 18626-1 | 18632-1 | -0.21 | 0.25 | 0.46 | 0.05 | 0.26 |
| | 18635-1 | 18624-1 | -1.21 | -0.05 | 1.16 | -0.07 | 1.14 |
| | 18631-1 | 18660-1 | 0.71 | -0.57 | 1.28 | -0.09 | 0.8 |
| | 17124-1 | 18631-1 | 0.26 | 1.57 | 1.31 | 0.42 | 0.16 |
| | 18631-1 | 18624-1 | 0.92 | 2.68 | 1.76 | 2.17 | 1.25 |
| | 18626-1 | 18660-1 | 0.17 | -2.74 | 2.91 | -2.08 | 2.25 |
| | | | | | MUE 0.94 | | MUE 0.95 |

| | Code ligand A | Code ligand B | Exp ΔΔG | Crystal Structure ΔΔG | Crystal Structure Absolute Error | $AF2_{30}$ ΔΔG | $AF2_{30}$ Absolute Error |
|---|---|---|---|---|---|---|---|
| MCL1 | 67 | 58 | -1.83 | -1.94 | 0.11 | -1.97 | 0.14 |
| | 52 | 60 | 0.31 | 0.19 | 0.12 | -0.26 | 0.57 |
| | 65 | 67 | 0.83 | 0.62 | 0.21 | 0.98 | 0.15 |
| | 63 | 60 | 0.15 | 0.9 | 0.75 | 1.25 | 1.1 |
| | 67 | 63 | -1.48 | -0.72 | 0.76 | -1.7 | 0.22 |
| | 56 | 35 | 0.45 | 2.93 | 2.48 | 1.2 | 0.75 |
| | 41 | 35 | -1.68 | 0.98 | 2.66 | -0.38 | 1.3 |
| | 35 | 53 | -1.15 | -4.03 | 2.88 | -2.15 | 1 |
| | 67 | 31 | -0.34 | 2.97 | 3.31 | 0.14 | 0.48 |
| | 67 | 35 | -1.23 | 2.43 | 3.66 | 0.98 | 2.21 |
| | | | | | MUE 1.69 | | MUE 0.79 |

| | Code ligand A | Code ligand B | Exp ΔΔG | Crystal Structure ΔΔG | Crystal Structure Absolute Error | $AF2_{30}$ ΔΔG | $AF2_{30}$ Absolute Error |
|---|---|---|---|---|---|---|---|
| P38 | p38a_2z | p38a_2y | 0.58 | 0.54 | 0.04 | -0.02 | 0.6 |
| | p38a_2z | p38a_3flw | -0.32 | -0.58 | 0.26 | -0.16 | 0.16 |
| | p38a_2aa | p38a_3flw | -1.41 | -1.96 | 0.55 | -0.06 | 1.35 |
| | p38a_2z | p38a_2aa | 1.09 | 1.67 | 0.58 | -0.1 | 1.19 |
| | p38a_2v | p38a_2y | -0.8 | -1.86 | 1.06 | -0.4 | 0.4 |
| | p38a_2v | p38a_2bb | -0.09 | -0.74 | 0.65 | -0.33 | 0.24 |
| | p38a_2z | p38a_3flq | 0.43 | 1.57 | 1.14 | 1.08 | 0.65 |
| | p38a_2z | p38a_3fmk | -1.47 | -2.93 | 1.46 | -3.02 | 1.55 |
| | p38a_2v | p38a_3fmk | -2.86 | -4.97 | 2.11 | -4.22 | 1.36 |
| | p38a_2aa | p38a_2bb | 0.2 | 2.51 | 2.31 | 0.95 | 0.75 |
| | | | | | MUE 1.02 | | MUE 0.83 |

| | Code ligand A | Code ligand B | Exp ΔΔG | Crystal Structure ΔΔG | Crystal Structure Absolute Error | $AF2_{30}$ ΔΔG | $AF2_{30}$ Absolute Error |
|---|---|---|---|---|---|---|---|
| PTP1B | 23467 | 23468 | -0.41 | -0.5 | 0.09 | 0.06 | 0.47 |
| | 23473 | 20669(2qbr) | -0.22 | -0.09 | 0.13 | 0.11 | 0.33 |
| | 23467 | 23473 | -1.05 | -1.21 | 0.16 | -0.77 | 0.28 |
| | 20669(2qbr) | 23472 | -0.04 | -0.23 | 0.19 | -0.91 | 0.87 |
| | 23467 | 23469 | -0.38 | -0.63 | 0.25 | 0.16 | 0.54 |
| | 23471 | 23468 | 0 | 0.94 | 0.94 | -0.47 | 0.47 |
| | 20670(2qbs) | 23482 | -0.92 | 0.06 | 0.98 | 0.14 | 1.06 |
| | 23471 | 23466 | -0.1 | 1.07 | 1.17 | 0.9 | 1 |
| | 23467 | 23475 | -1.38 | -2.68 | 1.3 | -0.6 | 0.78 |
| | 20670(2qbs) | 23466 | 1.24 | 3.24 | 2 | 1.02 | 0.22 |
| | | | | | MUE 0.72 | | MUE 0.60 |

| | Code ligand A | Code ligand B | Exp ΔΔG | Crystal Structure ΔΔG | Crystal Structure Absolute Error | $AF2_{30}$ ΔΔG | $AF2_{30}$ Absolute Error |
|---|---|---|---|---|---|---|---|
| Thrombin | 1b | 1a | 0.98 | 1.06 | 0.08 | 1.43 | 0.45 |
| | 1a | 5 | -0.1 | 0.08 | 0.18 | 0.79 | 0.89 |
| | 3a | 1b | -0.14 | -0.35 | 0.21 | -0.75 | 0.61 |
| | 1b | 7a | 0.24 | -0.09 | 0.33 | 0.85 | 0.61 |
| | 1b | 1c | -0.1 | -0.45 | 0.35 | -0.73 | 0.63 |
| | 3a | 1d | 0.07 | -0.83 | 0.9 | -2.56 | 2.63 |
| | 1a | 3b | -0.38 | 0.69 | 1.07 | 0.13 | 0.51 |
| | 1d | 1a | 0.77 | 2.42 | 1.65 | 2.63 | 1.86 |
| | 1d | 6e | -0.66 | 1.08 | 1.74 | 1.47 | 2.13 |
| | 1d | 5 | 0.67 | 3.02 | 2.35 | 3.03 | 2.36 |
| | | | | | MUE 0.89 | | MUE 1.27 |

| | Code ligand A | Code ligand B | Exp ΔΔG | Crystal Structure ΔΔG | Crystal Structure Absolute Error | $AF2_{30}$ ΔΔG | $AF2_{30}$ Absolute Error |
|---|---|---|---|---|---|---|---|
| Tyk2 | ejm_31 | ejm_45 | -0.02 | 0.01 | 0.03 | 0.08 | 0.1 |
| | ejm_45 | ejm_42 | -0.22 | -0.28 | 0.06 | -0.2 | 0.02 |
| | ejm_44 | ejm_42 | -2.36 | -2.27 | 0.09 | -3.25 | 0.89 |
| | jmc_23 | ejm_46 | 0.39 | 0.6 | 0.21 | -0.14 | 0.53 |
| | jmc_23 | jmc_30 | 0.76 | 0.54 | 0.22 | 0.74 | 0.02 |
| | ejm_31 | ejm_46 | -1.77 | -0.75 | 1.02 | -1.07 | 0.7 |
| | jmc_28 | jmc_30 | 0.04 | -0.52 | 0.56 | -0.49 | 0.53 |
| | ejm_47 | ejm_31 | 0.16 | -0.87 | 1.03 | -0.13 | 0.29 |
| | ejm_31 | jmc_28 | -1.44 | -0.39 | 1.05 | -0.2 | 1.24 |
| | ejm_42 | ejm_55 | 0.57 | -0.84 | 1.41 | -0.27 | 0.84 |
| | jmc_23 | ejm_55 | 2.49 | 0.83 | 1.66 | 0.53 | 1.96 |
| | | | | | MUE 0.67 | | MUE 0.70 |

Table S2: FEP+ results for general drug targets as reported in the original Wang et al[1].

| | Code ligand A | Code ligand B | Exp ΔΔG | Crystal Structure ΔΔG | Crystal Structure Absolute Error | $AF2_{30}$ ΔΔG | $AF2_{30}$ Absolute Error |
|---|---|---|---|---|---|---|---|
| A2A | 11 | 25a | 0.25 | 0.53 | 0.28 | 0.01 | 0.24 |
| | 11 | 25b | -1.15 | -0.34 | 0.81 | -0.88 | 0.27 |
| | 11 | 25c | -1.56 | -0.56 | 1 | -0.83 | 0.73 |
| | 11 | 25f | 0.35 | 0.23 | 0.12 | -0.66 | 1.01 |
| | 11 | 32 | 0.06 | 1.24 | 1.18 | -0.92 | 0.98 |
| | 11 | 41 | 0.56 | 3.08 | 2.52 | -1.35 | 1.91 |
| | 25a | 25f | 0.1 | -0.37 | 0.47 | -0.48 | 0.58 |
| | 25a | 32 | -0.18 | 1.32 | 1.5 | -1.08 | 0.9 |
| | 25b | 25c | -0.41 | -0.28 | 0.13 | -0.1 | 0.31 |
| | 25b | 25f | 1.51 | 0.62 | 0.89 | 0.49 | 1.02 |
| | 25d | 11 | 1.36 | 0.69 | 0.67 | 0.19 | 1.17 |

| | | | | | | |
|---|---|---|---|---|---|---|
| 25d | 25a | 1.6 | 1.85 | 0.25 | 0.9 | 0.7 |
| 25e | 11 | -1.39 | -0.88 | 0.51 | 0.69 | 2.08 |
| 25e | 25a | -1.14 | -0.79 | 0.35 | 0.33 | 1.47 |
| 25e | 25f | -1.04 | -0.76 | 0.28 | -0.37 | 0.67 |
| 25f | 25c | -1.92 | -1.25 | 0.67 | -0.48 | 1.44 |
| 41 | 25a | -0.32 | -0.88 | 0.56 | 1.33 | 1.65 |
| 41 | 32 | -0.5 | -0.5 | 0 | 0.29 | 0.79 |
| | | | | MUE 0.98 | | MUE 1.00 |

| | Code ligand A | Code ligand B | Exp ΔΔG | Crystal Structure ΔΔG | Crystal Structure Absolute Error | $AF2_{30}$ ΔΔG | $AF2_{30}$ Absolute Error |
|---|---|---|---|---|---|---|---|
| | 9 | 16 | -0.55 | 0.17 | 0.72 | 1.86 | 2.41 |
| | 9 | 17 | -0.04 | -0.64 | 0.6 | 1.11 | 1.15 |
| | 9 | 18 | 0.1 | -0.21 | 0.31 | 1.47 | 1.37 |
| B1AR | 9 | 14 | -1.09 | 0.32 | 1.41 | 1.26 | 2.35 |
| | 9 | 19 | -1.77 | 1.34 | 3.11 | -2.47 | 0.7 |
| | 9 | 15 | -0.35 | 1.32 | 1.67 | 2.18 | 2.53 |
| | 9 | 12 | -1.64 | -0.09 | 1.55 | -0.89 | 0.75 |
| | 9 | 13 | -1.5 | 0.75 | 2.25 | 1.14 | 2.64 |
| | 16 | 17 | 0.5 | -0.03 | 0.53 | -1.86 | 2.36 |
| | 17 | 18 | 0.14 | 0.01 | 0.13 | 0.58 | 0.44 |

| | | | | | | |
|---|---|---|---|---|---|---|
| 18 | 14 | -1.19 | 0.79 | 1.98 | -0.2 | 0.99 |
| 19 | 15 | 1.42 | 2.59 | 1.17 | 4.18 | 2.76 |
| 12 | 15 | 1.28 | 1.74 | 0.46 | 1.66 | 0.38 |
| 12 | 13 | 0.14 | 1 | 0.86 | 1.69 | 1.55 |
| 16 | 13 | -0.95 | -0.31 | 0.64 | -1.06 | 0.11 |
| | | | | MUE 1.16 | | MUE 1.50 |

Table S3: FEP+ results for GPCR targets as reported in Lenselink et al.[2]

| | Code ligand A | Code ligand B | Exp ΔΔG | Crystal Structure ΔΔG | Crystal Structure Absolute Error | $AF2_{30}$ ΔΔG | $AF2_{30}$ Absolute Error |
|---|---|---|---|---|---|---|---|
| JAK2 | 2_3E63 | 15 | 0.53 | 0.60 | 0.07 | -0.75 | 1.28 |
| | 2_3E63 | 13_3E64 | -1.79 | -2.00 | 0.21 | -2.10 | 0.31 |
| | 9 | 10 | 1.22 | 1.43 | 0.21 | -0.13 | 1.35 |
| | 5 | 15 | 0.41 | -0.01 | 0.42 | -0.42 | 0.83 |
| | 5 | 10 | 0.65 | 0.13 | 0.52 | 1.15 | 0.50 |
| | 12 | 14 | -0.35 | -0.94 | 0.59 | -0.40 | 0.05 |
| | 10 | 13_3E64 | -2.55 | -3.61 | 1.06 | -1.28 | 1.27 |
| | 21 | 13_3E64 | -1.36 | 0.14 | 1.50 | 0.54 | 1.90 |
| | 21 | 14 | 0.55 | 2.50 | 1.95 | 1.93 | 1.38 |
| | 12 | 13_3E64 | -2.26 | -4.80 | 2.54 | -1.49 | 0.77 |
| | | | | | MUE 0.91 | | MUE 0.96 |

| | Code ligand A | Code ligand B | Exp ΔΔG | Crystal Structure ΔΔG | Crystal Structure Absolute Error | $AF2_{30}$ ΔΔG | $AF2_{30}$ Absolute Error |
|---|---|---|---|---|---|---|---|
| HSP90 | 2 | 8 | -1.1 | -0.87 | 0.23 | 0.24 | 1.34 |
| | 9 | 11 | -0.76 | -1.25 | 0.49 | -0.7 | 0.06 |
| | 9 | 2 | 0.43 | 1.1 | 0.67 | 0.85 | 0.42 |
| | 12 | 10 | 0.28 | -0.65 | 0.93 | -0.73 | 1.01 |
| | 11 | 8 | 0.09 | 1.21 | 1.12 | 0.64 | 0.55 |
| | 11 | 12 | -0.47 | 0.9 | 1.37 | -0.65 | 0.18 |
| | 3 | 2 | 1.38 | -0.35 | 1.73 | 0.97 | 0.41 |
| | 12 | 14 | -0.13 | -2.81 | 2.68 | -3.36 | 3.23 |
| | 12 | 18 | -0.37 | -3.9 | 3.53 | -1.74 | 1.37 |
| | 18 | 10 | 0.65 | 5.74 | 5.09 | 1.04 | 0.39 |

| | | |
|---|---|---|
| | MUE 1.78 | MUE 0.90 |

Table S4: FEP+ results for fragments as reported in Steinbrecher et al.[3]

| | Code ligand A | Code ligand B | Exp ΔΔG | $AF2_{30}$ ΔΔG | $AF2_{30}$ Absolute Error |
|---|---|---|---|---|---|
| CDK2 (Shao et al., 2013) | 12b | 12a | -5.04 | -1.85 | 3.19 |
| | 12j | 12a | -2.23 | -1.19 | 1.04 |
| | 12j | 1b | -2.73 | -1.41 | 1.32 |
| | 12l | 12a | -1.93 | -0.62 | 1.31 |
| | 1b | 12a | -0.82 | -0.57 | 0.25 |
| | 1b | 12b | 4.22 | 1.34 | 2.88 |
| | 1b | 12l | 1.11 | 0.87 | 0.24 |
| | | | | | MUE 1.46 |

| | Code ligand A | Code ligand B | Exp ΔΔG | $AF2_{30}$ ΔΔG | $AF2_{30}$ Absolute Error |
|---|---|---|---|---|---|
| CDK9 (Shao et al., 2013) | 12b | 12a | -2.99 | -2.48 | 0.51 |
| | 12j | 12a | -0.09 | -1.68 | 1.59 |
| | 12j | 1b | -0.74 | -4.21 | 3.47 |
| | 12l | 12a | -0.77 | -1.14 | 0.37 |
| | 1b | 12a | 0.65 | 2.01 | 1.36 |
| | 1b | 12b | 3.64 | 2.91 | 0.73 |
| | 1b | 12l | 1.42 | 3.52 | 2.10 |
| | | | | | MUE 1.45 |

| | Code ligand A | Code ligand B | Exp ΔΔG | $AF2_{30}$ ΔΔG | $AF2_{30}$ Absolute Error |
|---|---|---|---|---|---|
| CDK2 (Blake et al., 2016) | 12 | 14 | 0.17 | 0.06 | 0.11 |
| | 12 | 18 | 0.58 | -1.01 | 1.59 |
| | 12 | 27 | 0.85 | -1.34 | 2.19 |
| | 16 | 12 | 0.71 | 1.18 | 0.47 |
| | 16 | 14 | 0.88 | 0.82 | 0.06 |
| | 18 | 27 | 0.27 | 0.10 | 0.17 |
| | 27 | 14 | -0.68 | 0.42 | 1.10 |
| | | | | | MUE 0.81 |

| | Code ligand A | Code ligand B | Exp ΔΔG | $AF2_{30}$ ΔΔG | $AF2_{30}$ Absolute Error |
|---|---|---|---|---|---|
| ERK2 (Blake et al., 2016) | 12 | 14 | 0.16 | -0.30 | 0.46 |
| | 12 | 18 | 2.19 | -0.75 | 2.94 |
| | 12 | 27 | 2.62 | -0.40 | 3.02 |
| | 16 | 12 | -0.31 | 0.48 | 0.79 |
| | 16 | 14 | -0.15 | 0.60 | 0.75 |
| | 18 | 27 | 0.43 | 0.52 | 0.09 |
| | 27 | 14 | -2.46 | 0.26 | 2.72 |
| | | | | | MUE 1.54 |

Table S5: FEP+ results in the context of specificity calculations as reported in Albanese et al.[4]

| | Code ligand A | Code ligand B | Exp DDG | Crystal Structure DDG (OPLS4) | Crystal Structure Absolute Error |
|---|---|---|---|---|---|
| MCL1 | 67 | 58 | -1.83 | -1.22 | 0.61 |
| | 52 | 60 | 0.31 | 0.35 | 0.04 |
| | 65 | 67 | 0.83 | 0.86 | 0.03 |
| | 63 | 60 | 0.15 | 0.56 | 0.41 |
| | 67 | 63 | -1.48 | -0.13 | 1.35 |
| | 56 | 35 | 0.45 | 1.51 | 1.06 |
| | 41 | 35 | -1.68 | -0.87 | 0.81 |
| | 35 | 53 | -1.15 | -2.05 | 0.9 |
| | 67 | 31 | -0.34 | 1.79 | 2.13 |
| | 67 | 35 | -1.23 | 1.96 | 3.19 |
| | | | | | MUE = 1.05 |

| | Code ligand A | Code ligand B | Exp DDG | Crystal Structure DDG (OPLS4) | Crystal Structure Absolute Error |
|---|---|---|---|---|---|
| P38 | p38a_2z | p38a_2y | 0.58 | 1.04 | 0.46 |
| | p38a_2z | p38a_3flw | -0.32 | -0.32 | 0 |
| | p38a_2aa | p38a_3flw | -1.41 | -1.29 | 0.12 |
| | p38a_2z | p38a_2aa | 1.09 | 0.48 | 0.61 |
| | p38a_2v | p38a_2y | -0.8 | -0.97 | 0.17 |
| | p38a_2v | p38a_2bb | -0.09 | -0.96 | 0.87 |
| | p38a_2z | p38a_3flq | 0.43 | -0.48 | 0.91 |
| | p38a_2z | p38a_3fmk | -1.47 | -2.59 | 1.12 |
| | p38a_2v | p38a_3fmk | -2.86 | -4.67 | 1.81 |
| | p38a_2aa | p38a_2bb | 0.2 | 0.79 | 0.59 |
| | | | | | MUE = 0.67 |

Table S5: FEP+ results for MCL1 and P38 using crystal structures and the latest version of the OPLS forcefield (OPLS4)

**Supplementary Figures**

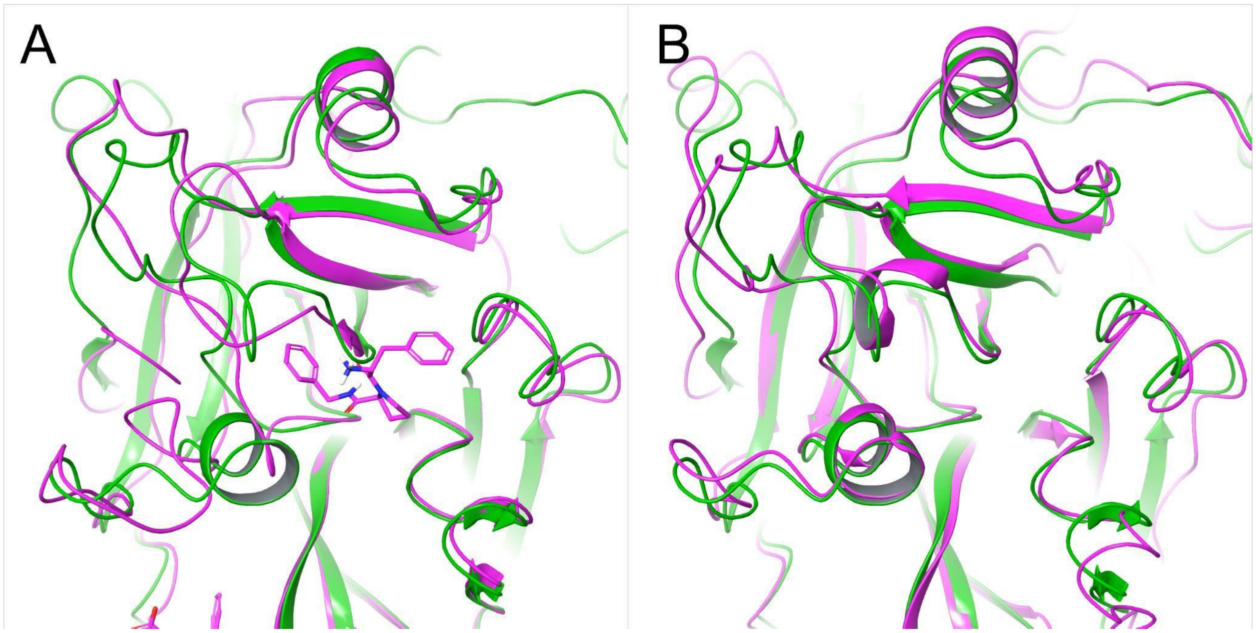


**Figure S1**. AF2 model (green) superimposed on Thrombin reference structures 2ZFF (A) and 6C2W (B)

A2A (P29274)

| | | | |
|---|---|---|---|
| 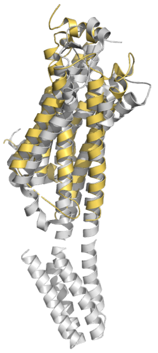 | 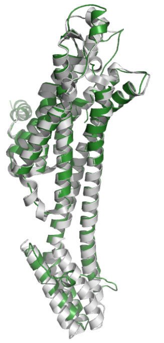 | 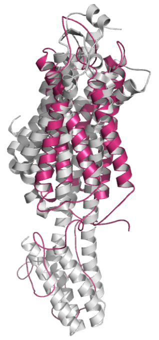 | 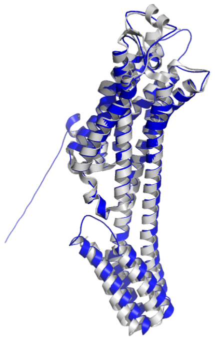 |
| 10.09 | 2.22 | 13.60 | 2.09 |

B1AR (P08588)

| | | | |
|---|---|---|---|
| 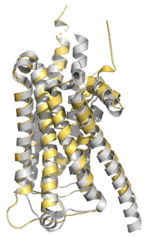 | 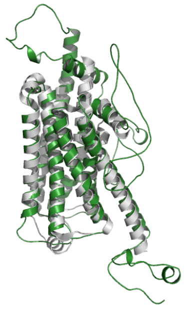 | 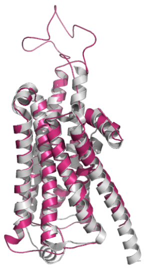 | 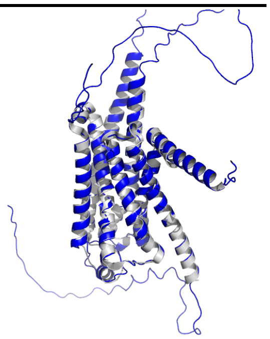 |
| 3.51 | 3.36 | 3.81 | 1.01 |

BACE (P56817)

| | | | |
|---|---|---|---|
| 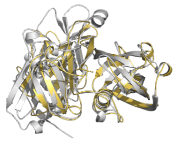 | 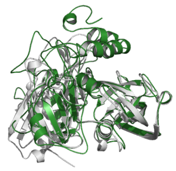 | 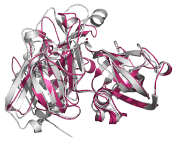 | 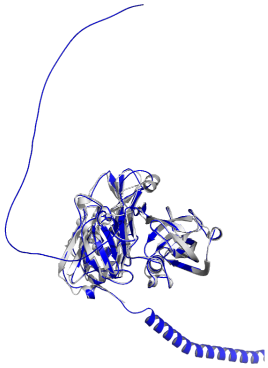 |
| 6.04 | 9.12 | 4.94 | 1.90 |
| CDK2 (P24941) | | | |
| 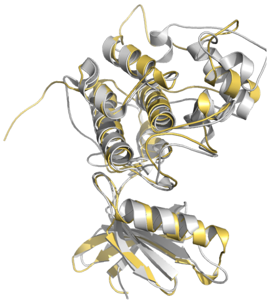 | 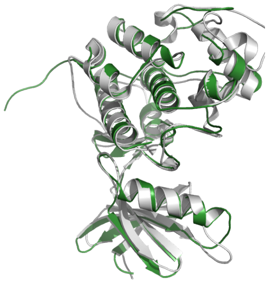 | 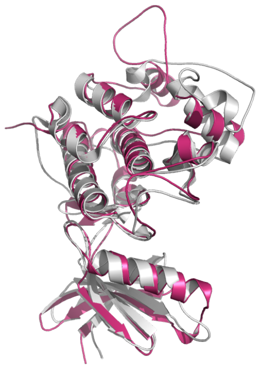 | 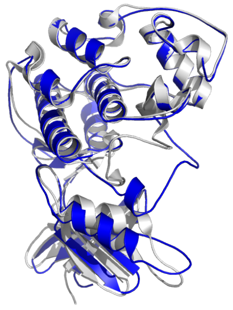 |
| 6.53 | 5.81 | 7.35 | 3.66 |
| CDK9 (P50750) | | | |

| | | | |
|---|---|---|---|
| 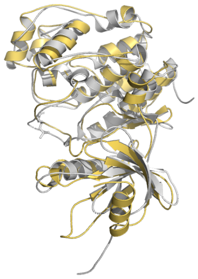 | 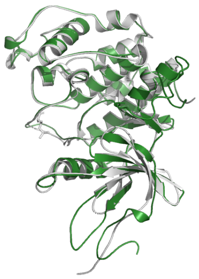 | 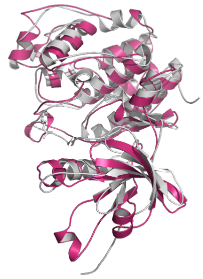 | 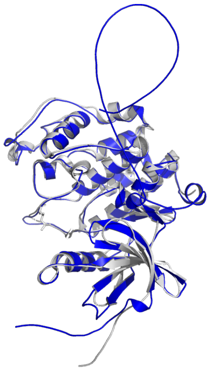 |
| 5.47 | 1.59 | 4.88 | 1.64 |
| ERK2 (P28482) | | | |
| 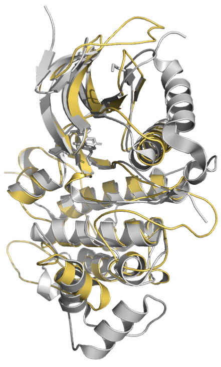 | 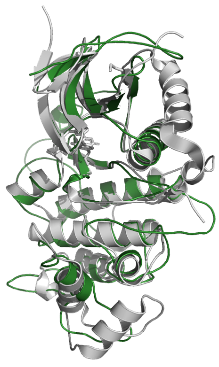 | 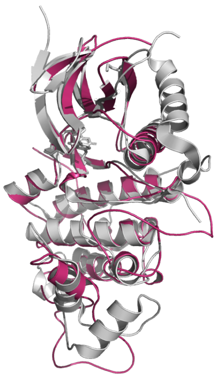 | 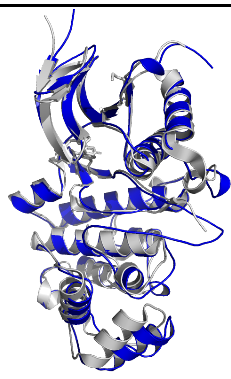 |
| 12.59 | 14.85 | 7.40 | 1.47 |
| HSP90 (P07900) | | | |

| | | | |
|---|---|---|---|
| 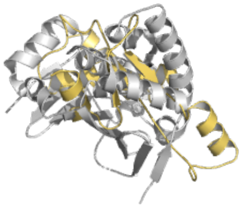 | 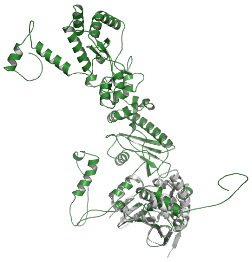 | 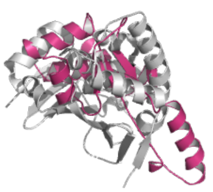 | 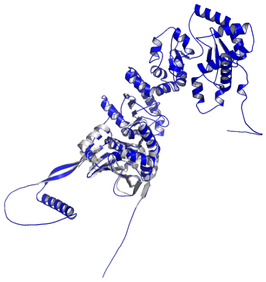 |
| 20.15 | 9.04 | 7.61 | 0.88 |
| JAK2 (O60674) | | | |
| 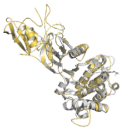 | 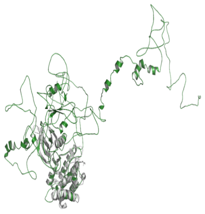 | 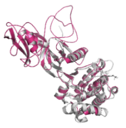 | 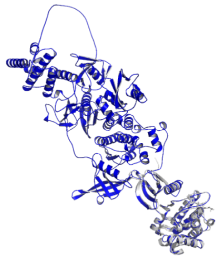 |
| 7.07 | 13.56 | 5.01 | 0.98 |
| JNK1 (P45983) | | | |

| | | | |
|---|---|---|---|
| 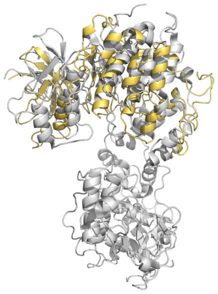 | 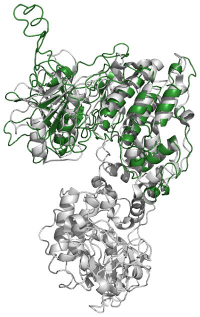 | 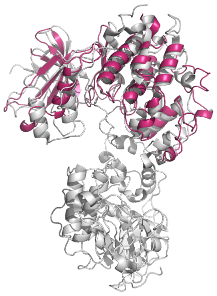 | 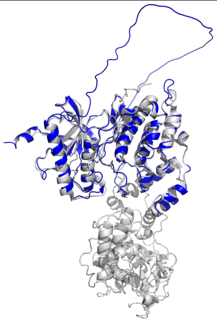 |
| 12.61 | 10.71 | 7.79 | 1.40 |
| MCL1 (Q07820) | | | |
| 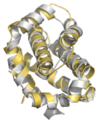 | 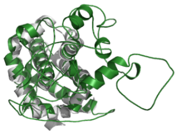 | 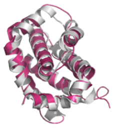 | 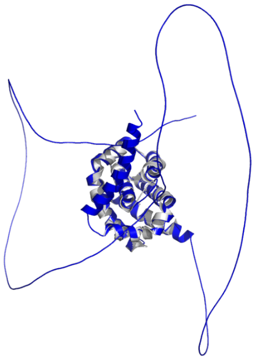 |
| 3.03 | 4.19 | 4.45 | 1.12 |
| p38 (Q16539) | | | |

| | | | |
|---|---|---|---|
| 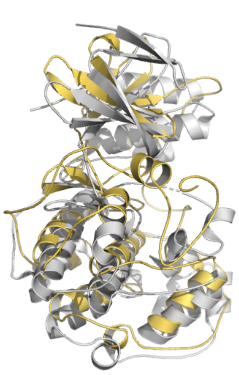 | 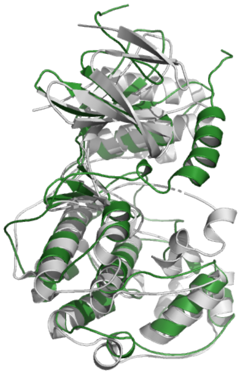 | 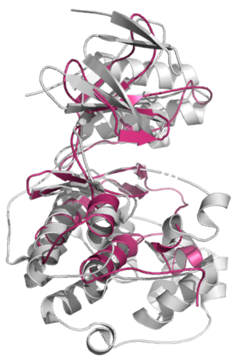 | 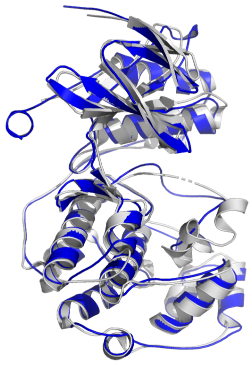 |
| 13.44 | 11.27 | 6.39 | 2.85 |
| PTP1B (P18031) | | | |
| 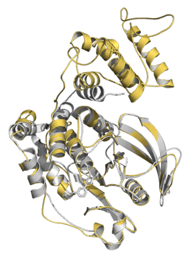 | 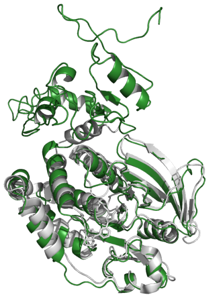 | 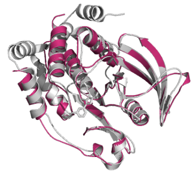 | 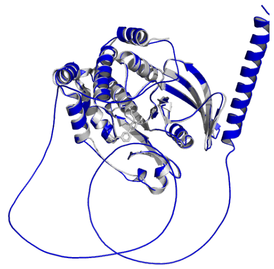 |
| 7.41 | 4.56 | 2.00 | 0.75 |
| Thrombin (P00734) | | | |

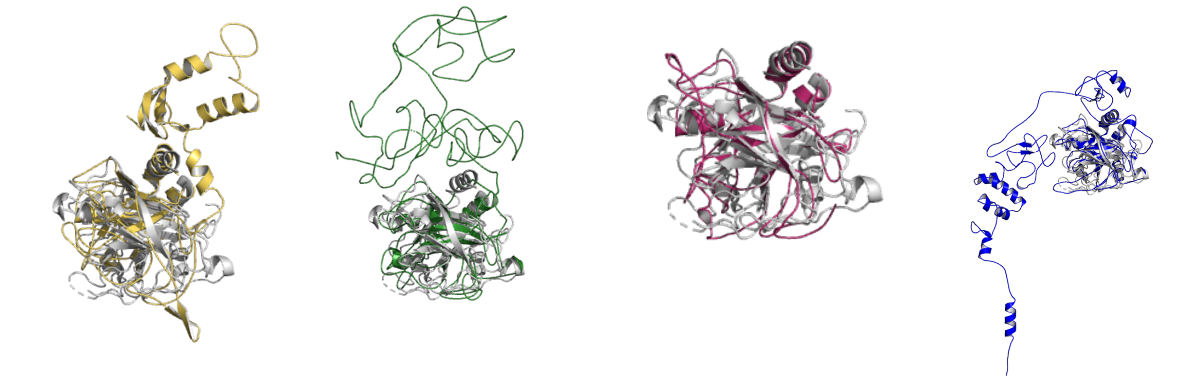

6.59 4.84 3.79 1.53

Tyk2 (P29597)

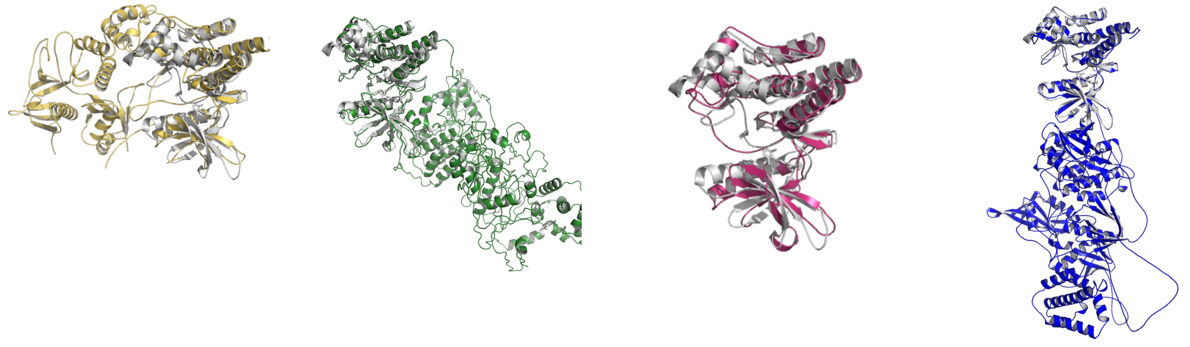

6.45 2.85 4.90 1.13

**Figure S2**. Comparison of the reference crystal structure (light gray) with the homology models obtained by Prime (yellow), I-TASSER (green) and SwissModel (red) and $AF2_{30}$ (blue) using templates with a maximum identity threshold of 30%. Global Cα-RMSD (in Å) is indicated for each system.

### Supplementary Information References


1. Wang, L. *et al*. Accurate and reliable prediction of relative ligand binding potency in prospective drug discovery by way of a modern free-energy calculation protocol and force field. *J. Am. Chem. Soc.* 137, 2695–2703 (2015).

2. Lenselink, E. B. *et al*. Predicting binding affinities for GPCR ligands using free-energy perturbation. *ACS Omega* 1, 293–304 (2016).

3. Steinbrecher, T. B. *et al*. Accurate binding free energy predictions in fragment optimization. *J. Chem. Inf. Model.* 55, 2411–2420 (2015).

4. Albanese, S. K. *et al*. Is structure-based drug design ready for selectivity optimization? *J. Chem. Inf. Model.* 60, 6211–6227 (2020).
